# Iron Chelator-Mediated Anoxic Biotransformation of Lignin by Novel sp., *Tolumonas lignolytica* BRL6-1

**DOI:** 10.1101/2020.05.14.095802

**Authors:** Gina Chaput, Andrew Billings, Lani DeDiego, Roberto Orellana, Joshua N Adkins, Carrie Nicora, Rosalie Chu, Blake Simmons, Kristen M. DeAngelis

## Abstract

Lignin is the second most abundant carbon polymer on earth and despite having more fuel value than cellulose, it currently is considered a waste byproduct in many industrial lignocellulose applications. Valorization of lignin relies on effective and green methods of delignification, with a growing interest in the use of microbes. Here we investigate the physiology and lignin biotransformation mechanisms of the novel facultative anaerobic bacterium, *Tolumonas lignolytica* BRL6-1, under anoxic conditions. Physiological and biochemical changes were compared between cells grown anaerobically in either lignin-amended or unamended conditions. In the presence of lignin, BRL6-1 had a higher biomass and shorter lag phase compared to unamended conditions, and 14% of the proteins determined to be significantly higher in abundance by log_2_ fold-change of 2 or greater were related to Fe(II) transport in early exponential phase. Ferrozine assays of the supernatant (<10 kDa fraction) confirmed that Fe(III) was bound to lignin and reduced to Fe(II) only in the presence of BRL6-1, suggesting redox activity by the cells. LC-MS/MS analysis of the secretome showed an extra band at 20 kDa in lignin-amended conditions. Protein sequencing of this band identified a protein of unknown function with homology to enzymes in the radical SAM superfamily. Expression of this protein in lignin-amended conditions suggests its role in radical formation. From our findings, we suggest that BRL6-1 is using a protein in the radical SAM superfamily to interact with the Fe(III) bound to lignin and reducing it to Fe(II) for cellular use, increasing BRL6-1 yield under lignin-amended conditions. This interaction potentially generates organic free radicals and causes a radical cascade which could modify and depolymerize lignin. Further research should clarify the extent to which this mechanism is similar to previously described aerobic chelator-mediated Fenton chemistry or radical producing lignolytic enzymes, such as lignin peroxidases, but under anoxic conditions.

## Introduction

The industrial processing of lignocellulosic material produces 5×10^6^ metric tons of lignin annually worldwide (1). Lignin is the largest renewable source of aromatics that can be used for products such as flavors, fragrances, dyes, and other valuable secondary metabolites (2,3). However, it is considered an “untapped” resource due to the recalcitrant nature of the polymer, making it difficult to separate and process for valuable downstream products (4,5).

Investigation of microbial mediated processes for the depolymerization of lignin have focused predominantly on aerobic fungi and bacteria (4,6–9). Under oxic conditions, enzymes such as laccases and peroxidases produce oxidants that diffuse into and reduce the lignin complex (9,10), causing bond scission reactions between lignin subunits. For organisms that lack lignolytic enzymes such as brown-rot fungi as well as bacteria like *Pantoea ananatis* Sd-1 and *Cupriavidus basilensis* B-8, chelator-mediated Fenton chemistry (CMF) is used to depolymerize lignin (11–14). In this mechanism, the microorganism produces an iron reducer molecule, a chelator molecule, and H_2_O_2_. Once the chelator binds to Fe(III) in the environment, it then reacts with the iron reducer molecule to reduce Fe(III) to Fe(II). Fe(II) then reacts with H_2_O_2_ to create •OH radicals. Similarly to oxidants formed by laccases and peroxidases, the •OH radicals disrupt the lignin structure, causing bond scissions of subunits (14). Both lignolytic enzyme and chelator mediated lignin depolymerization are promising for industries that rely lignocellulosic feedstocks (8). For example, the use of Fenton chemistry for lignocellulosic processing has been studied using the aerobic bacterium, *C. basilensis* B-8 grown on rice straw, showing a synergistic relationship in lignin depolymerization and cellulose yield (13). However, limitations to these processes hinder them from being competitive on the market. Both aerobic fungi and bacteria require constant aeration and mixing, making it very costly to maintain the cultures (15). Mass production of fungal or bacterial lignolytic enzymes are also not possible due to lacking a method of recycling the enzymes after one use, low substrate specificity, and low redox potential (16). For example, both laccases and manganese-dependent peroxidases cannot degrade 80-90% of lignin due to the presence of non-phenolic structures (17).

Though originally thought to be impossible 30 years ago (18), anaerobic bacteria could offer a solution to issues presented for aerobic microorganisms. For example, based on genome analysis, *Klebsiella sp*. strain BRL6-2 is hypothesized to use lignin as an electron acceptor for energy production (19). Support for this mechanism is also seen with humic substances, which are lignin rich (20), acting as extracellular electron acceptors for bacteria in sediments and anoxic waters (21). *Enterobacter lignolyticus* SCF1 has been studied comparing the growth in lignin-amended and unamended conditions under anaerobic conditions. RNAseq analysis suggested various enzymes that may be responsible for lignin depolymerization, including alcohol dehydrogenases (22,23). However, the exact mechanism has yet to be elucidated. By identifying additional anaerobic bacteria capable of degrading lignin, mechanisms and their regulation can be uncovered and further utilized for lignin depolymerization and valorization applications.

*Tolumonas lignolytica* BRL6-1 is a novel, facultative anaerobic bacterium that was isolated from the El Yunque experimental forest, Puerto Rico, on lignin as the sole carbon (C) source under anoxic conditions (24). Previously, BRL6-1 has demonstrated to have a shorter lag phase and a higher biomass in the presence of lignin (24). However, the mechanism of lignin modification and how it benefits cell growth is not well understood. We hypothesize that when grown anaerobically in the presence of lignin, BRL6-1 produces an extracellular protein that acts as both iron chelator and redox agent. This protein potentially generates organic free radicals and causes a radical cascade that modifies and depolymerizes lignin.

## Materials and methods

### Culturing *Tolumonas lignolytica* BRL6-1

To study lignin modification under anoxic conditions, *Tolumonas lignolytica* BRL6-1 was grown in 0.04% D-glucose as the primary C source amended or unamended with 0.1% alkali lignin, low sulfonate (Sigma Aldrich, CAS Number 8068-05-1). Cultures grew on modified CCMA media consisting of (per liter) 2.25 g NaCl, 0.5 g NH_4_Cl, 0.227 g KH_2_PO_4_, 0.348 g K_2_erPO_4_, 5 mg MgSO_4_•7H_2_O, 2.5 mg CaCl_2_•2H_2_O, 0.01 mL SL-10 trace elements, and 0.01 mL Thauer’s vitamins (25–27). The D-glucose concentration was 0.2% for the ferrozine assays, Arnow assays, proteome and secretome analysis, described in more detail below. Cultures grew at 30°C anaerobically in triplicate and uninoculated bottles served as abiotic controls. Iron amended cultures had an additional 38 ppb Fe(II) added to the media as FeCl_2_•4H_2_O.

To study the physiological response of BRL6-1 in the presence of lignin, growth in lignin-amended medium, lignin unamended medium, and lignin unamended medium supplemented with the additional Fe (II) were monitored by measuring cell concentration by adsorption (OD_600_). Bacterial growth curves were analyzed with gcFit function via *grofit* package in R (28). Calculated average lag phase, maximum growth rate (μ Max), and maximum cell growth (A) were based on the Gompertz Model.

### Proteomic Analysis for Cell Pellet and Secretome

To identify proteins differentially expressed during lignin-amended growth, biomass was collected at early and late logarithmic growth phase from cultures grown in the presence or absence of lignin. Briefly described, proteins were extracted from cell pellet fractions using methanol/chloroform and analyzed with LC-MS/MS. Raw mass spectrometry data were searched with MS-GF+ against NCBI RefSeq *Tolumonas* sp. BRL6-1 database (October 2014 version) in addition to bovine/por cine trypsin and other common contaminants such as keratin sequences (3164 total sequences). Searching parameters required tryptic digestion of at least one of the peptide ends (partially tryptic), <10 ppm peptide mass tolerance and methionine oxidation as variable modification. The identified MS/MS spectra were filtered with an MS-GF+ score of 1e^-09^ resulting in ≤ 1.0% false discovery rate (FDR) at the protein level. The count of spectra attributed to each individual protein within each experimental condition was used as a proxy for relative quantitative values.

Supernatant fractions from late logarithmic growth phase were collected to identify differentially expressed proteins in the secretome during lignin-amended growth. Cell-free supernatant was generated by collecting 100 mL cultures at the end of late logarithmic growth phase and vacuum filtrated with a 0.45 μm filter. Cell-free supernatant was centrifuged using a 10 kDa filter & the >10 kDa fraction was further concentrated using trichloroacetic acid (TCA) precipitation (41). Samples were run on a 15% SDS-PAGE gel and silver stained. Bands of interest from both lignin-amended and unamended samples were cut out at 20 kDa, 37 kDa, and 50 kDa. Using an in-gel tryptic digest kit (Thermo Fisher, Catalog #89871), samples were prepared as described by the manufacturer for LC-MS/MS analysis. LC-MS/MS analysis was completed by the Mass Spectrometry Center at University of Massachusetts Amherst. Raw mass spectrometry data was search with MS/GF+ against the NCBI RefSeq *T. lignolytica* BRL6-1 database (2016 version).

### Statistical Analysis of Proteomic Data

Spectral counts from the cell pellet proteomics between lignin-amended and unamended conditions were compared using msms.edgeR function via *msms.Tests* package in R (29). The post-test effect size filter of msms.edgeR deemed proteins differentially expressed if proteins had p-values <0.05, absolute values of log_2_ fold-change >1 or < -1, and total spectral counts >2 across biological replicates (29).

### Ferrozine and Arnow Assays

In order to confirm that iron was bound to lignin, Fe(II) and Fe(III) concentrations were measured with ferrozine assays as described with modifications (30). Supernatant from lignin-amended and unamended cultures were harvested by removing 15 mL aliquots of culture from serum bottles under anoxic conditions during lag phase, late logarithmic growth phase, and mid-stationary growth phase. Samples were filtered under anoxic conditions through a 0.45 µm filter to remove biomass and then ultrafiltrated with a 10 kDa Amicon filter. Filtrate (<10 kDa) was tested in triplicate for total iron. Briefly, in a 96-well plate under anoxic conditions, 225 µL of sample with 15 µL 1 M ascorbic acid were added, followed by 60 µL of 50 mg ferrozine mL^-1^ and 500 mM potassium acetate buffer, per 5.5. Plates were wrapped in tin foil and incubated for 135 min at 37°C before being measured at 562 nm with a plate spectrophotometer. A separate set of plates had ascorbic acid substituted with water to calculate Fe(II) in the media. For both ferrozine assays a standard curve of Fe(II) was completed as well as samples taken from the abiotic cultures to act as controls.

To determine if BRL6-1 is producing catechol-like siderophores known as catecholates, Arnow assays were completed as described (31). This assay was performed using the same fractions as the ferrozine assays. Briefly, 1 mL of a <10 kDa fraction sample was combined with 1 mL 0.5 M HCl, 1 mL nitrite-molybdate reagent, and 1 mL 1 M NaOH. Reactions were incubated for 5 min before being diluted 5-fold with water in a 96-well plate and light absorbance read at 510 nm. Samples from the biotic replicates as well as abiotic controls were completed in triplicate. A standard curve was produced using 3,4-dihydroxybenzoate (31).

### Inductively Coupled Plasma (ICP) Spectroscopy of Kraft Alkali Lignin Substrate

One gram of alkali lignin, low sulfonate (Sigma Aldrich, CAS Number 8068-05-1) was sent for analysis in triplicate to the University of Massachusetts Amherst Soil and Plant Nutrient Testing Laboratory. Lignin was acid wet digested in nitric acid, hydrochloric acid, and hydrogen peroxide in a block digester and measured with ICP Spectroscopy to determine the total P, K, Ca, Mg, Zn, Cu, Mn, Fe, and B.

## Results and discussion

### Physiology and global expression changes in response to lignin

To determine the role of lignin in the anaerobic metabolism and growth of *T. lignolytica* BRL6-1, we compared cultures grown with glucose that were either amended or unamended with lignin. Lignin-amended cultures had shorter lag phase (5.0 ± 0.6 hrs) compared to cells grown in unamended conditions (11.0 ± 4 hrs, p-value =0.03). Lignin-amended cultures also had higher yields (0.140 ± 0.013 OD_600_) compared to cells grown in unamended conditions (0.124 ± 0.003 OD_600,_ p-value = 0.04). Maximum growth rate in unamended conditions was not significantly different from lignin-amended conditions (0.039 ± 0.03 OD_600_ hr^-1^ compared to 0.03 ± 0.008 OD_600_ hr^-1^. Our growth results support previous findings that under lignin-amended conditions, BRL6-1 fitness improves, experiencing a shorter lag phase and achieving a higher final biomass when lignin is amended (24).

To explain this change in growth, we originally hypothesized that lignin may serve as a secondary C source as well as a potential energy source (24). To explore this hypothesis, we compared protein production between lignin-amended conditions to unamended. EdgeR analysis uses the negative binomial distribution to detect proteins that are differentially abundant between the two growth conditions (29). The analysis resulted in a total of 41 proteins with significantly higher abundance and 101 lower abundance relative to unamended conditions in early exponential phase and a total of 9 proteins with significantly higher abundance and 9 proteins lower abundance in late exponential phase (**Fig 1**).

**Fig 1.**
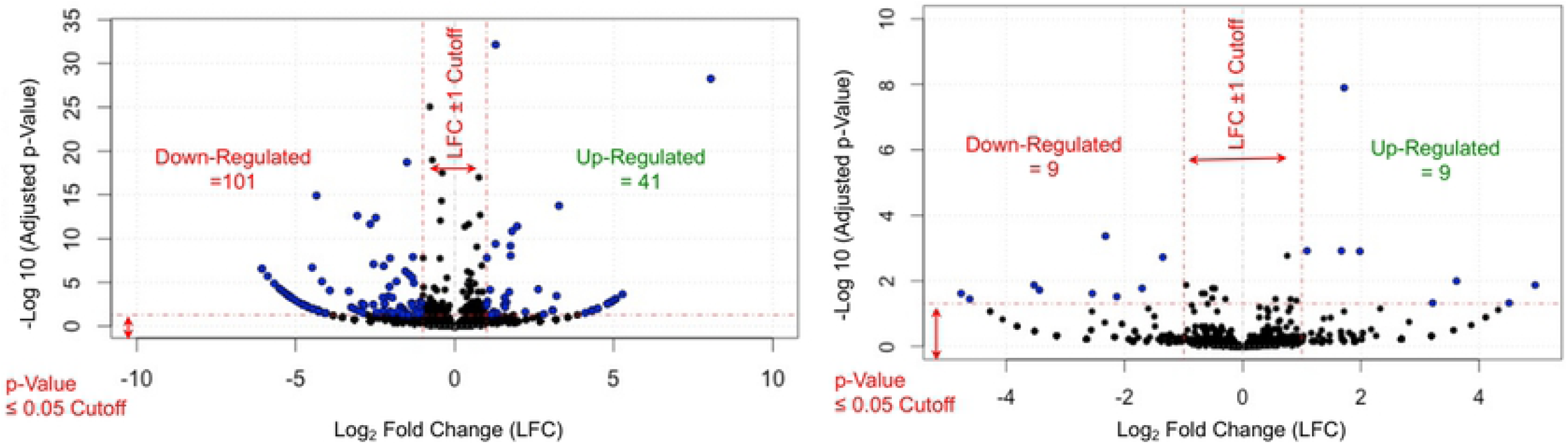
Proteomic analysis results of *T. lignolytica* BRL6-1 grown in lignin-amended versus unamended conditions. Blue dots represent significant change in protein abundance in lignin-amended conditions (p-value < 0.05) whereas black dots denote represent proteins that did not change between lignin-amended and unamended conditions.

### Evidence of proton-relay system-related enzyme for lignin degradation

From our edgeR analysis results, we first investigated the roles of proteins that had significantly higher abundance in both early and late exponential phase in lignin-amended cultures relative to unamended. This subset of proteins included the protein with the highest abundance listed in lignin-amended conditions in both phases, which was annotated as a carboxymuconolactone decarboxylase (CMD) family protein (Pfam 02627). The CMD family of proteins is associated with aromatic degradation in aerobic bacteria, via the protocatechuate branch of the β-ketoadipate pathway, as well as antioxidant defense via peroxidase activity (32,33). For our protein, we identified an alkylhydroperoxidase (AhpD) domain and CXXC motif (log_2_ fold-change of 8 and 7, respectively) (**Fig 2A**). Based on the CXXC motif, it is thought that the protein detected in BRL6-1 has AhpD-like activity (32). AhpD is part of an antoxidant defense system that forms a complex with peroxiredoxin, AhpC. The function of AhpD is to restore the enzyme activity of AhpC via reduction (34). Looking further into the genome, BRL6-1 contains a gene annotated as AhpF, which is an alternative alkyl hydroperoxide reductase to AhpD as seen in *Salmonella typhimurium* (63.5% sequence identity with NCBI BLASTp) (35). Based on this information, AhpC likely forms a complex with AhpF. To support this, BPROM program analysis, which predicts bacterial promoter sites, placed AhpC and AhpF downstream of the same predicted promoter in the BRL6-1 genome (36). Additionally, our edgeR analysis determined AhpCF protein complex expression was not significantly different between lignin-amended and unamended conditions. Therefore, this up-regulated AhpD-like protein in the presence of lignin could be serving another role with its reducing activity instead of restoring AhpC activity.

**Fig 2.**
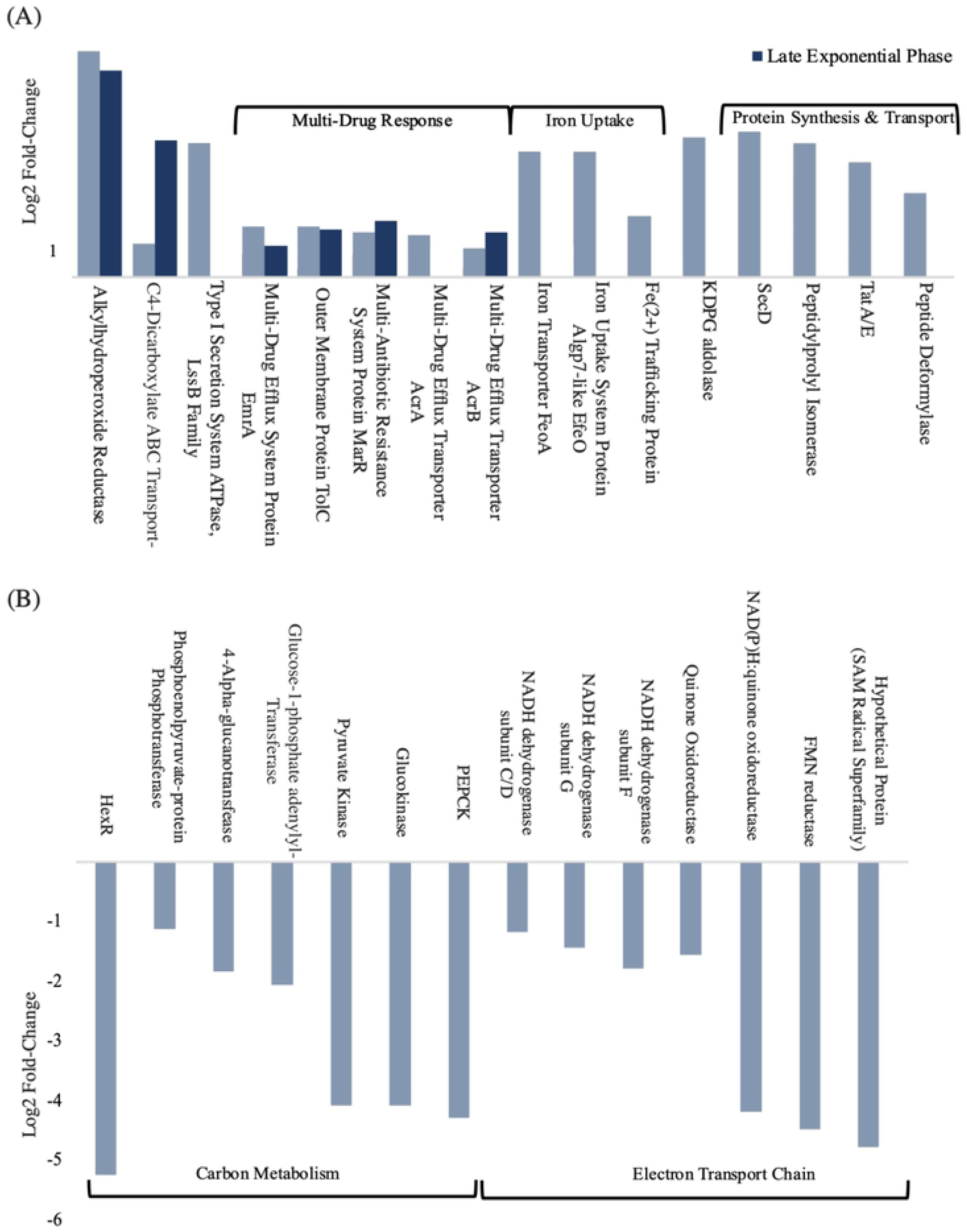
Proteins with significant change in abundance (log_2_ fold change, p-value < 0.05) related to carbon metabolism and energy production by *T. lignolytica* BRL6-1 under lignin-amended conditions relative to unamended. (A) Proteins of BRL6-1 with significantly high and (B) low abundance under lignin-amended conditions compared to lignin unamended (p-value < 0.05). Abbreviations are the following: phosphoenolpyruvate carboxykinase (PEPCK); 2-keto-3-deoxy-6-phosphogluconate (KDPG).

The mechanism that AhpD uses to reduce its substrates is a proton relay system (37), which was originally described as protons being shuttled from the active site of a protein to bulk solvent molecules (38). This mechanism has been observed previously in the lignin degrading enzyme, LigL, found in *Sphingomonas paucimobilis* SYK-6. LigL catalyzes stereospecific oxidation of the benzylic alcohol as the first degradation step of lignin-derivative, (αS, βR)-GGE (39,40). The proton relay mechanism has also been described for p-Cresol methylhydroxylase (PCMer) in *Pseudomonas* species to degrade phenol p-cresol as well as p-hydroxybenzyl alcohol (37). Therefore, this AhpD-like protein could be reducing lignin-derived compounds in the cell using this mechanism. In support of cellular aromatic compound uptake, a C4-dicarboxylate ABC transporter protein in both early and late exponential phase (log_2_ fold-change of 1.2 and 4.9, respectively). This transporter has an 87% sequence identity to transporter, DctA, in *Pseudomonas chlororaphis* O6 that was found to be essential for benzoate uptake (41).

### Lignin-amended cultures shit in carbon metabolism and energy production

With evidence that BRL6-1 is potentially transporting benzoate compounds into the cell and reducing them via AphD, we then compared changes in protein abundance relating to carbon (C) metabolism in the presence of lignin. In lignin-amended conditions, the HexR transcriptional factor was significantly lower in BRL6-1 by a log_2_ fold-change of -5 during early logarithmic growth phase (**Fig 2B**). HexR is a global transcription factor utilized for fine-tuning the carbohydrate catabolic pathways to adapt to variable C availability in the environment (42–44). It has been previously shown that HexR is present in several groups of Proteobacteria and regulates a rather complex set of operons (44). Leyn and colleagues (2011) deleted *hexR*, which resulted in the de-repression of transcription of the central glycolytic genes as well as the activation of genes involved in gluconeogenesis. In contrast to what would be expected, our results show that BRL6-1’s glucokinase and pyruvate kinase, which are responsible for the first and last step of glycolysis, respectively, were lower in abundance by a log_2_ fold-change of -4, suggesting that the presence of lignin lead to the repression of glycolysis (**Fig 2**). This evidence is also supported by the fact that phosphoenolpyruvate carboxykinase (PEPCK), the rate limiting enzyme for gluconeogenesis, was also lower in abundance in lignin-amended growth conditions by a log2 fold-change of -4, indicating that PEP was not being funneled into gluconeogenesis, whereas a KDPG aldolase was significantly higher in abundance by a log_2_ fold-change of 5 (**Fig 2**). These findings suggest that in the presence of lignin, there is a higher conversion of KDPG to pyruvate by the KDGP aldolase. Very dynamic transcription regulation patterns on C metabolism pathways have been previously reported during metabolization of lignin of *E. lignolyticus* SCF1, another facultative anaerobic Gammaproteobacteria isolated from tropical rainforest soils (22).

Despite the observed shifts in C metabolism-related proteins described above, enzymes related to aromatic metabolism that were previously annotated in the genome did not significantly change in abundance in the presence of lignin relative to the unamended conditions (24). This lack in change may be due to the fact that either no monomers are being cleaved from lignin in the presence of BRL6-1 or that lignin was depolymerized into high molecular weight polymers and not directly consumed by BRL6-1.

To determine if lignin had a role in energy production, we searched for enzymes that were related to the electron transport chain that had differential abundance in lignin-amended conditions compared to unamended. During early exponential phase, 6% of all significantly lower abundance proteins (log_2_ fold-change >1, p-value <0.05) in the presence of lignin were NADH dehydrogenase subunits (log_2_ fold-change between -1.2 to -4) and one flavin mononucleotide (log_2_ fold-change of -4) (**Fig 2B**).

### Iron redox by *T. lignolytica* BRL6-1 in the presence of lignin and hypothesized mechanisms

Of the proteins that were significantly higher in abundance by a log_2_ fold-change >2 in the presence of lignin during early exponential phase, 14% were related to Fe(II) uptake (**Fig 2A**). We hypothesized that BRL6-1 could be obtaining energy using iron redox and that lignin could play a role due to its strong affinity for iron (45). The complex between lignin and iron makes iron more soluble in the environment but not necessarily more bio-available for cellular use (46). This considered, BRL6-1 may have a mechanism that is disrupting the lignin-iron association. By doing so, BRL6-1 could obtain both iron and a potential C source faster than cells in unamended conditions, explaining the ability of BRL6-1 to exit lag phase more quickly in the presence of lignin (47).

To further investigate the relationship between lignin, iron, and BRL6-1 fitness, we first asked if lignin was contributing to a higher iron concentration aside from the SL-10 minerals we added to the media. ICP Spectrophotometry showed that the lignin substrate contained 38 ppb iron (data not shown). To test whether the 38 ppb Fe was benefitting BRL6-1 fitness, we completed Fe addition growth curve experiments and monitored BRL6-1 growth. Compared to cells grown in lignin unamended conditions reported above, lignin unamended conditions with 38 ppb iron addition were not significantly different in lag phase, growth rate, or biomass yield: 10.5 ±5 hrs, 0.072 ±0.03 OD_600_ per hr, 0.124 ±0.007 OD_600_, respectively. These results suggest that additional iron alone was not enough to benefit BRL6-1 growth.

To test if lignin has a strong affinity for iron in our system, we completed a ferrozine assay for <10 kDa fractions of supernatant from lag phase, late logarithmic growth phase, and mid-stationary growth phase. We expected that if the iron was binding to lignin in the media, we should see less bioavailable iron in the supernatant of lignin-amended compared to lignin unamended conditions. In lignin-amended conditions, Fe(III) was not detectable in the <10 kDa fractions throughout the entire growth curve whereas 292 ppb Fe(III) was detected during lag phase in lignin unamended conditions. As bacterial biomass increased over time, Fe(II) accumulated in both conditions to similar concentrations (**Fig 3**). There was no change in Fe(IIMIII) concentrations in abiotic controls. This suggests that Fe(III) was bound to the lignin and was reduced to Fe(II) by BRL6-1.

**Figure 3.**
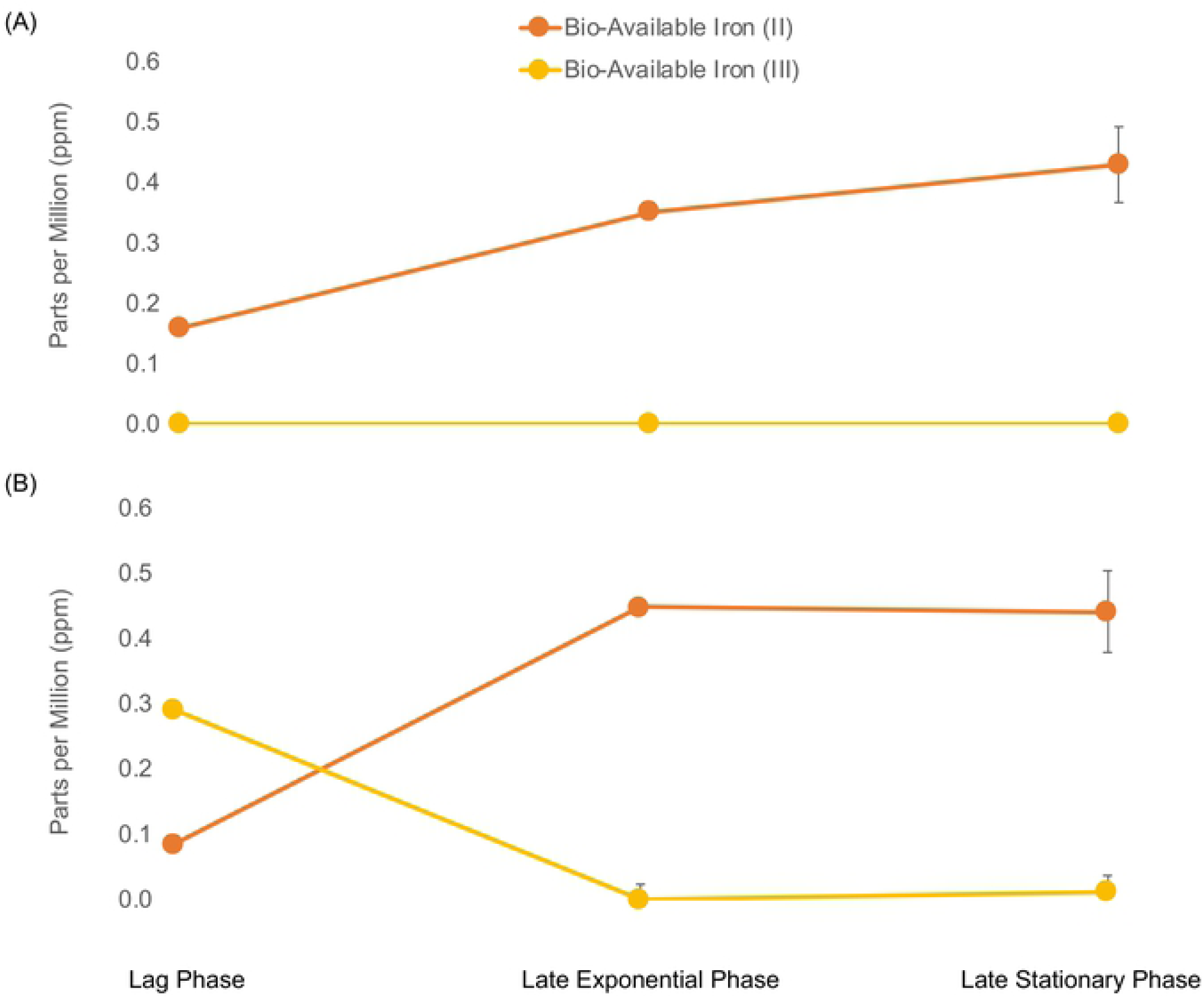
Bio-available Fe(II) (orange) and Fe(III) (yellow) concentrations in parts per million (ppm). Concentrations of Fe(II/III) when BRL6-1 was cultured in (A) lignin-amended and (B) in unamended conditions at lag phase, late exponential phase, and late stationary phase.

With evidence for Fe(III) bound to lignin and being reduced to Fe(II) based on ferrozine assays, we next wanted to determine the mechanism that BRL6-1 may use to accomplish this redox reaction. One potential explanation was the use of siderophores, which are organic molecules used by bacteria to chelate Fe(III) under iron-limiting conditions (48). The most common siderophore used by bacteria are catecholates, which rely on hydroxyl groups of the catechol rings to form the iron chelation center (31). To see if BRL6-1 produces this type of siderophore, we completed Arnow assays on <10 kDa supernatant fractions from lignin-amended and unamended cultures during lag phase, late logarithmic growth phase, and mid-stationary growth phase. Catecholate detection was seen only in lignin-amended conditions; however, there was no change in concentration of catechol over the course of the growth curve (**Fig 4**). Additionally, abiotic controls of the lignin-amended conditions had similar concentrations to biotic replicates. This is likely due to Arnow assays being non-specific between catecholates and compounds containing catechol, such as soluble lignin (49), making it difficult to differentiate sources as well as any small changes in concentration of such metabolites. BRL6-1 may also be producing other groups of siderophores such as hydroxamates or carboxylates (48), which would need to be detected with a Csáky assay or the use of phenolphthalein and sodium hydroxide, respectively (50,51). Despite these limitations, initial results suggest that siderophores are unlikely to be the main explanation for the Fe redox being detected by the ferrozine assay and instead another mechanism may be used by BRL6-1.

**Fig 4.**
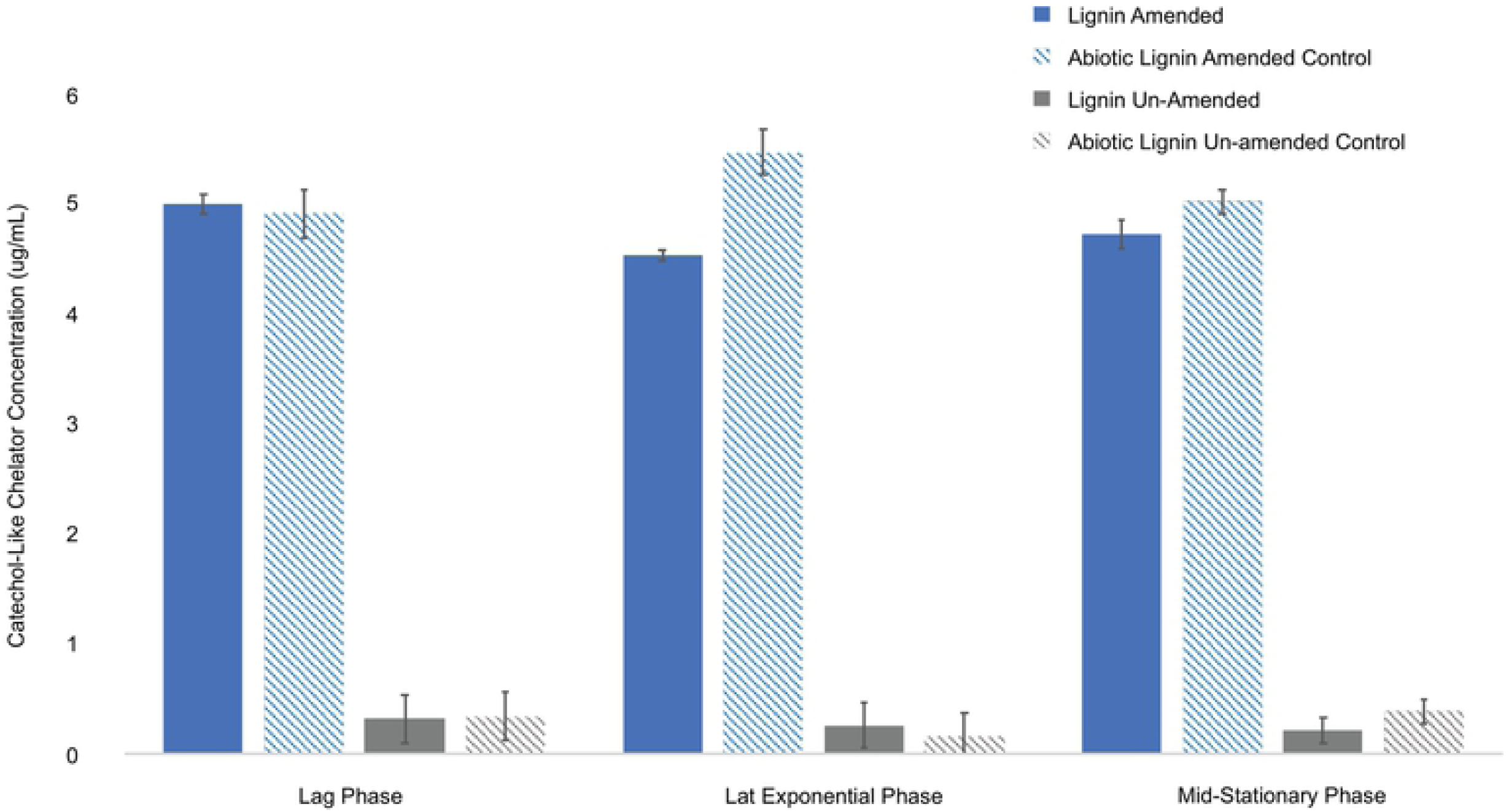
Arnow assay of catechol-like compounds present in supernatant. Catechol-like chelator concentrations (*µ*g/mL)at lag, late exponential, and mid-stationary phase of BRL6-1growth under lignin-amended (blue) and unamended conditions (gray). Abiotic controls are striped for both conditions.

We investigated to see if BRL6-1 secreted lignolytic enzymes or iron reducing proteins in the presence of lignin as seen for aerobic fungi and bacteria (11,12,14). Samples from late stationary phase were run on an SDS-PAGE and silver stained to detect differential banding between the two growth conditions. Differential banding was at 20kDa under lignin-amended conditions (**Fig. 5**). In addition, because BRL6-1 has had previously predicted peroxidases in its genome (24) and that lignin peroxidases are 35-48 kDa (52), we also were interested in the bands at 37 and 50kDa. Therefore, bands were cut out at 50 kDa, 37 kDa, and 20 kDa for both conditions to identify the proteins present by sequencing.

**Fig 5.**
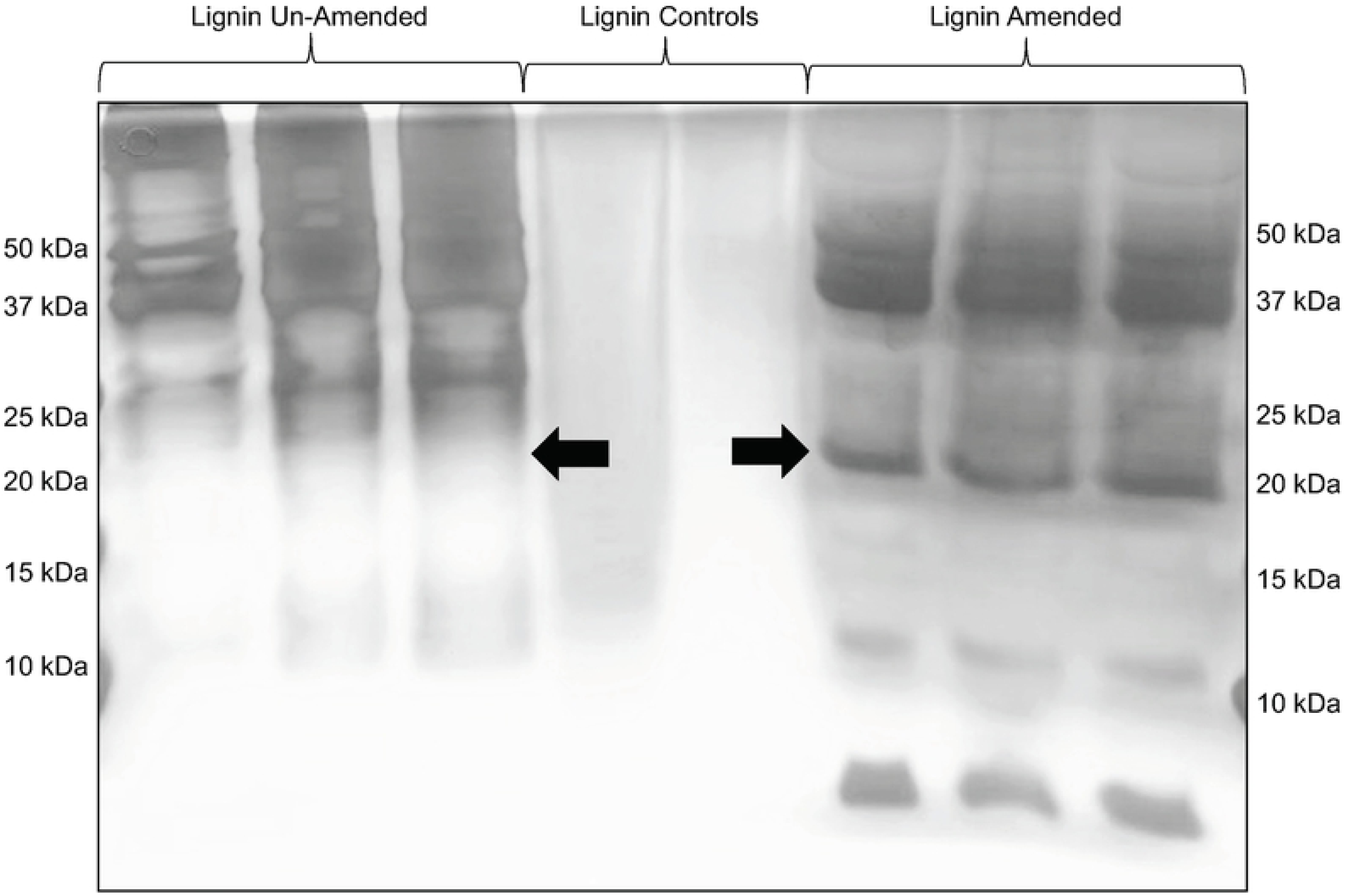
SDS-PAGE of *T. lignolytica* BRL6-1 secretome cultured in lignin-amended and unamended conditions. Arrows showing differential banding at 20kDa.

A protein originally annotated as hypothetical, WP_024871222.1 was detected in all three lignin-amended biological replicates with a predicted size of 20.8 kDa. There were no conserved domains detected in this protein, but Position-Specific Iterated (PSI) BLAST analysis of the protein identified homology to hypothetical protein from *Alteromonadales bacterium* BS08 (53% Identity; E-value 4e-61). BS08 was isolated from the gut of *Bankia setacea*, also known as the shipworm, that digests wood as a food source (53). Additionally, WP_024871222 had homology to enzymes in the radical SAM superfamily (37.7% Identity; E-value 0.06), a wide range of enzymes that support oxygen-independent alternatives to aerobic pathways (54). This may be relevant since *Tolumonas lignolytica* as other *Tolumonas* spp., are the only members of the *Aeromonadaieae* that are oxidase and catalase negative (24). Therefore, it is possible, that WP_024871222.1 has a role in lignin modification via radical formation.

## Conclusions

The presence of lignin has been previously shown to be beneficial for the growth of *T. lignolytica* BRL6-1 (24) with speculation that lignin was acting as a secondary C source or energy source. In our study, proteomic analysis revealed that 14% of the upregulated proteins by log_2_ fold-change of 2 or greater relating to Fe(II) transport in early exponential phase. Transient iron accumulation in *Salmonella enterica* serovar Typhimurium is required in order for the cells to come out of lag phase (47) and so it was possible that the up-regulation of iron enzymes might be due to BRL6-1 having the same iron requirement or relied on iron redox for energy (55). However, lignin has a strong affinity for iron (45) and therefore we would expect that iron is less bioavailable to the cells in lignin-amended conditions (46). Ferrozine assays of the <10kDa supernatant fractions confirmed that Fe(III) was bound to lignin, but it was reduced to Fe(II) when BRL6-1 was present, suggesting redox activity by the cells. To explain this redox activity, we hypothesized that BRL6-1 is producing a protein that acts as both an iron chelator and redox agent under anoxic conditions to obtain the iron bound to lignin. Secretome (extracellular enzyme) analysis coupled with LC-MS/MS identified the presence of a protein of unknown function but had homology to enzymes in the radical SAM superfamily, suggesting that it may have a role in radical formation in lignin-amended conditions.

While our work illustrates a potential molecular mechanism for anaerobic lignin modification by *T. lignolytica* BRL6-1, protein isolation and characterization are needed to confirm that this protein interacts with the Fe(III) bound to lignin and reduces it to Fe(II) for cellular use. Further analysis with electron paramagenetic resonance (EPR) as well as nuclear magnetic resonance (NMR) are necessary in order to confirm that organic radicals are being produced in the process of this enzyme obtaining iron and that these radicals alter the structure of lignin. By continuing research into the biochemical nature of microbial lignin transformations under anaerobic conditions, industries using lignocellulose as raw material will be one step closer to lignin valorization.

## Acknowledgements

Mass spectral data were obtained at the University of Massachusetts Mass Spectrometry Center. This work was part of the DOE Joint BioEnergy Institute (http://www.jbei.org) supported by the U. S. Department of Energy, Office of Science, Office of Biological and Environmental Research, through contract DE-AC02-05Cer11231 between Lawrence Berkeley National Laboratory and the U. S. Department of Energy. This research was performed using resources at the DOE Environmental Molecular Sciences Laboratory (grid.436923.9), which is an DOE Office of Science User Facilities sponsored by the Office of Biological and Environmental Research and operated under Contract No. DE-AC05-76RL01830. This publication was also developed under Assistance Agreement No. FP-91782301-0 awarded by the U.S. Environmental Protection Agency to Gina Chaput. It has not been formally reviewed by EPA. The views expressed in this document are solely those of Gina Chaput *et al*. and do not necessarily reflect those of the Agency. EPA does not endorse any products or commercial services mentioned in this publication.

